# Exploring the distribution of statistical feature parameters for natural sound textures

**DOI:** 10.1101/2020.08.28.271528

**Authors:** Ambika P. Mishra, Nicol S. Harper, Jan W.H. Schnupp

## Abstract

Sounds like “running water” and “buzzing bees” are classes of sounds which are a collective result of many similar acoustic events and are known as “sound textures”. Recent psychoacoustic study using sound textures by [1] reported that natural sounding textures can be synthesized from white noise by imposing statistical features such as marginals and correlations computed from the outputs of cochlear models responding to the textures. The outputs being the envelopes of bandpass filter responses, the ‘cochlear envelope’. This suggests that the perceptual qualities of many natural sounds derive directly from such statistical features, and raises the question of how these statistical features are distributed in the acoustic environment. To address this question, we collected a corpus of 200 sound textures from public online sources and analyzed the distributions of the textures’ marginal statistics (mean, variance, skew, and kurtosis), cross-frequency correlations and modulation power statistics. A principal component analysis of these parameters revealed a great deal of redundancy in the texture parameters. For example, just two marginal principal components, which can be thought of as measuring the sparseness or burstiness of a texture, capture as much as 66% of the variance of the 128 dimensional marginal parameter space, while the first two principal components of cochlear correlations capture as much as 90% of the variance in over 1000 correlation parameters. Knowledge of the statistical distributions documented here may help guide the choice of acoustic stimuli with high ecological validity in future research.

---

To address this question, we analyzed the marginal statistics (mean, variance, skew, and kurtosis), as well as cochlear envelope correlations and modulation power statistics of a sound corpus of 200 sound textures collected for this purpose. Using principal component analysis, we explored the distributions of these statistical parameters. Variabilities explained by the first two principal components of marginals, correlations and modulation power statistics are 65%, 90% and 73%, respectively.

As we progress in the first principal component space of marginal statistics, sounds which are naturally more “sparse” have higher “variance, skew and kurtosis” and with lower “mean” for the sound envelopes. The second principal component explains sounds which have lower envelope variance across frequency bands. The first principal component of correlation statistics distinguishes “highly correlated” to “poorly correlated” sounds across all frequency bands, whereas the second principal component can explain sounds which are “highly correlated “for lower frequency bands only. Sounds with low modulation power for low modulation frequency bands and high modulation power for high modulation frequency bands are captured by the first principal component of modulation power statistics whereas second principal component have the opposite trend to first principal component. This study further suggests that large “acoustic variability” of natural sound texture space can be compensated by significantly small “statistical variability”.

## Introduction

Be it buzzing bees, a flowing river, flocks of squawking birds or howling wind, the natural world is filled with a huge diversity of different sound textures, and humans have added further to that variety with all manner of traffic and machine noises. While this variety may at first glance seem limitless, it is nevertheless useful to ask whether all this variety of environmental sound can be captured in a more or less bounded, finite parameter space with knowable parameter distributions. If so, estimating the parameter distributions that characterize a large portion of the perceptual diversity of environmental sounds could be very useful, as it would allow us to ask how sound stimuli used in psychoacoustic or physiological studies of the auditory system relate to the types of sounds the auditory system actually encounters, and may have adapted to.

Here we describe our attempt to characterize these distributions by collecting and statistically analysing a large corpus of a class of natural sounds known as sound textures. We understand sound textures in the sense popularized by [1], as sounds that may have a lot of complexity, like for example the sound of waves breaking on a pebble beach, but which are nonetheless fully described by a finite set of stationary statistical parameters. Hence highly realistic exemplars of such sounds can be synthesized from scratch by morphing random noise samples to assume the spectral, modulation power and cross-frequency correlation structure characteristic of that type of texture. While textures defined in this way are fundamentally stochastic, and thereby exclude some important classes of sounds which are highly deterministic (such as highly regular rhythms) or rule based (such as a spoken sentence or a piece of music) they nevertheless cover a large proportion of the sounds encountered in natural and man-made environments, and the fact that they appear to be well characterized by a potentially large but finite number of stationarity statistical parameters makes the research questions we are pursuing here tractable.

Previous studies have identified a variety of parameters that are in principle suitable for characterizing sounds, including, for example, features like Mel-frequency cepstral coefficients (MFCC), band energy ratio, spectral flux and the wavelet subspace cepstrum. These have been nicely reviewed by [2], and are often used in applications such as sound event classification and computational auditory scene analysis (CASA). However, cepstral coefficients tend to look at relatively short time windows, so here we chose to use the auditory texture statistics by [1], which were inspired by the previous characterization of features used in visual texture discrimination research, [3], [4], [5].

[1] were able to show that natural sounding textures can be synthesized de novo by “shaping-noise” to impose the statistical features of the desired sound on random noise samples. Synthesized sounds from “white noise” by [1] were often easily identifiable as exemplars of a particular type of natural sound, and in many cases indistinguishable from a natural recording. The statistical parameters they adopted in their study was inspired by knowledge of the filtering of sounds known to be performed by the peripheral and central auditory systems. In their model, the input signal is band pass filtered into a range of frequency bands which mimics cochlear filtering. The amplitude envelope of the signal in each frequency band is extracted and cochlear transduction of sound is simulated by applying compressive nonlinearity to the amplitude envelopes (raising the envelope by a power of 0.3).

From the compressed cochlear envelopes, statistics such as mean, variance, skew, kurtosis and correlation between bands are computed for the amplitude distributions of these “cochlear envelopes”. The mean, variance, skew and kurtosis (also referred to as the first, second, third and fourth moment respectively) of the envelope amplitudes, will collectively be referred to as “marginal moments”, or “marginals” of the sound texture. In addition, the pairwise correlations between cochlear envelope amplitudes (“cochlear correlations”, for short) are computed. Previous studies by [5, 6] reported that both marginal moments and correlations are important features of visual textures, and the same is clearly true for auditory textures. Furthermore, the compressed cochlear envelopes are also passed through a second bank of band pass filters to measure the distribution of amplitude modulations (the “modulation power” statistics) and to compute correlations between modulation channels.

To appreciate how the types of statistical parameters extracted by the [1] model can distinguish types of sound textures, consider that some textures are “sparse”, exhibiting periods of relative silence with burst of energy of widely varying amplitude distribution (e.g. grains of hail bouncing on a tin roof), while others have a much more constant stream of sound (e.g. a high pressure jet of water rushing out of a faucet). Marginal moments will distinguish sparse from less sparse sounds easily, with sparse sounds having relatively greater variance, skew and kurtosis. Indeed, the usefulness of marginal moments in distinguishing natural sounds and images has been appreciated for a while [7, 8]. Similarly, modulation power statistics may be useful to distinguish “buzzing insects” sound textures, which have low modulation power at low frequencies but higher modulation power at higher frequency bands, from “waves on a beach” for which modulation power is relatively uniform across all modulation frequency bands.

In addition to cochlear marginals and modulation power distributions, [1] also examined the role of correlations, either between cochlear envelopes, or between modulation filters. Cochlear correlations (C) turned out to be a perceptually very powerful feature, discriminating, for example, the sound of applause, in which the ebb and flow of acoustic energy is highly correlated across many cochlear frequency channels, from “running water” type sounds, in which correlations between cochlear envelopes are small. Unlike cochlear correlations, modulation correlations are computed between outputs of modulation filter banks, and they come in two “flavors”, cross-modulation-frequency-band (C1) and within-modulation-frequency-band (C2). But unlike cochlear correlations, modulation correlations do not appear to play a less important role in auditory texture perception [1], and we shall not consider modulation correlations further in this study.

## 1 Materials and methods

To examine the statistical parameter space spanned by natural sound textures, we collected a corpus of natural sound recordings, computed their statistical parameters using the [1] framework, and subjected the resulting database of statistical parameters to dimensionality reduction by principal component analysis (PCA). This allowed us to identify parsimonious “principal feature axes” which explain a substantial portion of the variability among sound textures typically found in the environment, and to determine the ranges of parameter values that environmental sound textures typically occupy.

### 1.1 Sound collection

We collected 450 high quality raw sound samples from freesound, a freely available web resource [9]. After a preliminary inspection, we selected 200 sound samples which were deemed to be “texture like”. Sound clips with long duration of silence were excluded. All the sounds are of 48 kHz sample rate, and each clip is of 15s duration.

### 1.2 Statistical parameter extraction

Hence in this study, we explore the distribution of marginal moments, cochlear correlation, and modulation power over our corpus of natural sound textures. There are thus three aspects to our study of the statistical parameters of natural textures: a) Exploring the marginal moments. b) Exploring the correlation statistics. c) Exploring the modulation power statistics. The workflow of our exploration process for each statistical feature type is shown in Fig 1.

**Fig 1.**
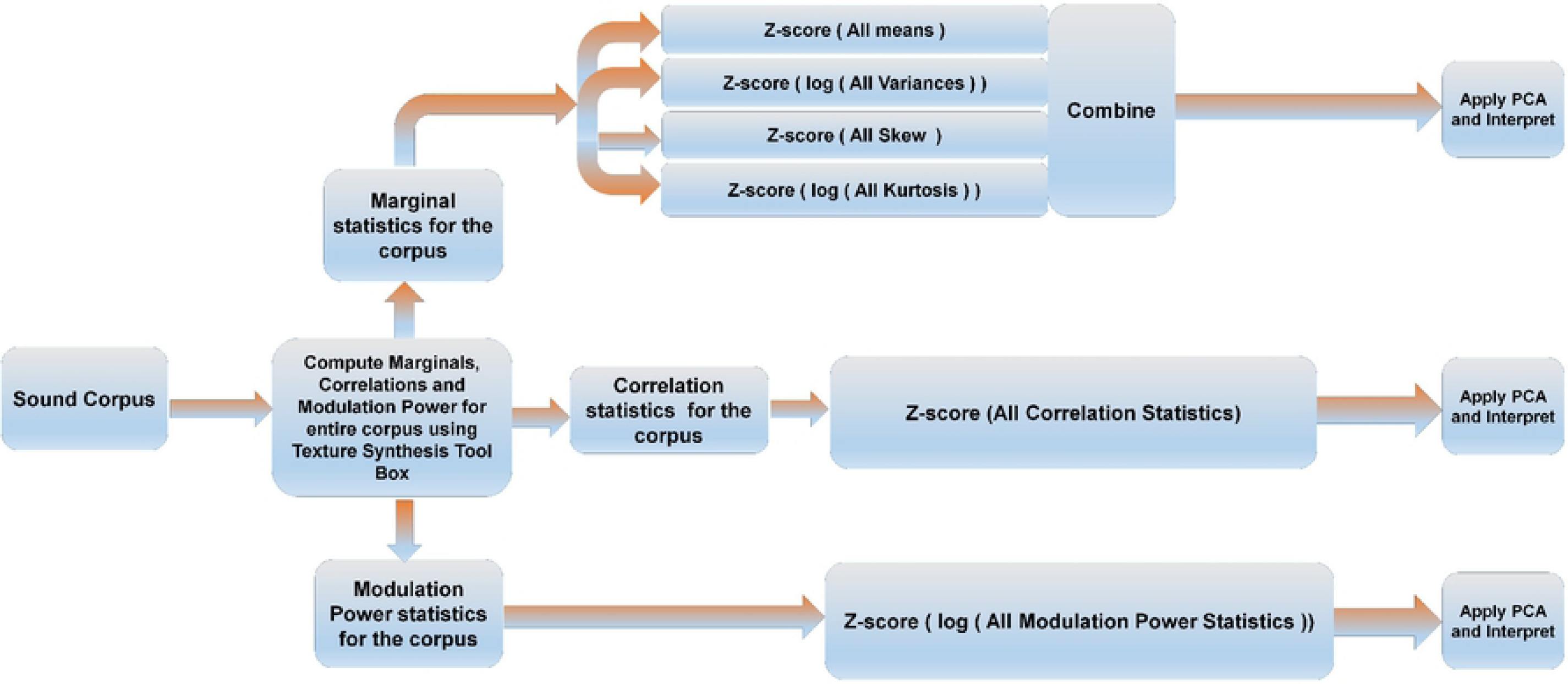
Work flow for exploring the statistical parameter space of a corpus of natural sounds. Using the sound synthesis tool box, marginal, correlation and modulation power statistics were computed for the entire corpus. Envelope variance and kurtosis, as well as modulation power parameters, are log transformed to make their distributions more symmetric around the mean. Each of these parameter sets is normalized and centered by z-scoring, and the z-scored parameter sets are subjected to PCA and interpreted.

The Sound Synthesis Toolbox V1.7 by [1] segregates each input sound into a number of “cochlear” frequency sub bands and computes the marginal statistics for each sub band. Here we have taken 32 cochlear filters with center frequencies equally spaced on an equivalent rectangular bandwidth (ERB) scale [10], spanning 80-20000 Hz. which is similar to the model by [1]. The output of each of the 32 cochlear filters undergoes envelope extraction and compression, and four “marginal moments”, mean, variance, skew and kurtosis, of the envelope values are computed, yielding 32×4=128 marginal parameters for each sound.

The Sound Synthesis Toolbox V1.7 also computes the pair-wise correlation between the cochlear envelopes, yielding 32×32=1024 correlation parameters (although these are somewhat redundant given that the correlation matrix is symmetric around the main diagonal). To compute the modulation power parameters, the output of each cochlear envelope is passed through another set of 20 “modulation” bandpass filters. The center frequencies of these modulation filters are equally spaced on a log scale from 0.5 to 200 Hz (same parameters to those used by [1]. Modulation power is then measured as the variance (mean sum of squares) of the output of each modulation filter, normalized by the variance of the respective cochlear envelope. For each sound, a total of 32 (cochlear channels) ×20 (modulation channels)=640 modulation power parameters are computed.

Thus, each sound in our corpus is described by a parameter set of 128 marginal values, 1024 correlation values and 640 modulation values, a very high-dimensional parameter space, but also one that is expected to be highly redundant, given that, for example, the marginal moments in adjacent frequency bands are bound to be highly correlated. To examine this redundancy, and to arrive at a low-dimensional parametrization of our sound corpus which would make it feasible to examine the ranges and distributions of statistical features that are common among environmental sounds, we subjected each of the parameter sets (marginals, correlations, modulations) to PCA. Prior to PCA, the raw parameter values underwent the following two pre-processing steps: Firstly, the distributions of the envelope variance and kurtosis parameters, as well as the modulation power parameters, were strongly positively skewed, as might be expected. They were therefore log-transformed to yield more symmetric and compact parameter distributions. Secondly, the distributions of means, log(variances), skew and log(kurtosis), as well as those of the correlations and the log(modulation power) values were normalized and centred by z-scoring, respectively. After these preprocessing steps, the matrices of 200(sound examples)×128 marginal parameters, 200×1024 correlations, and 200×640 modulation power values were independently analysed with PCA.

## 2 Results

The distributions of the original and transformed (pre-processed) statistical parameters computed for the corpus are shown in Fig 2A-2H. Fig 2A shows the distribution of the mean envelope amplitudes across frequency bands for the entire corpus. The distribution of variances of the envelope amplitudes is shown in Fig 2C, and the distributions of skew and kurtosis are depicted in Fig 2E and 2G respectively. The distributions of the raw variance and kurtosis parameters in particular are quite asymmetric, with a noticeable positive skew. This asymmetry is reduced after log transformation and z-scoring, as can be seen in Fig 2D and 2H.

**Fig 2.**
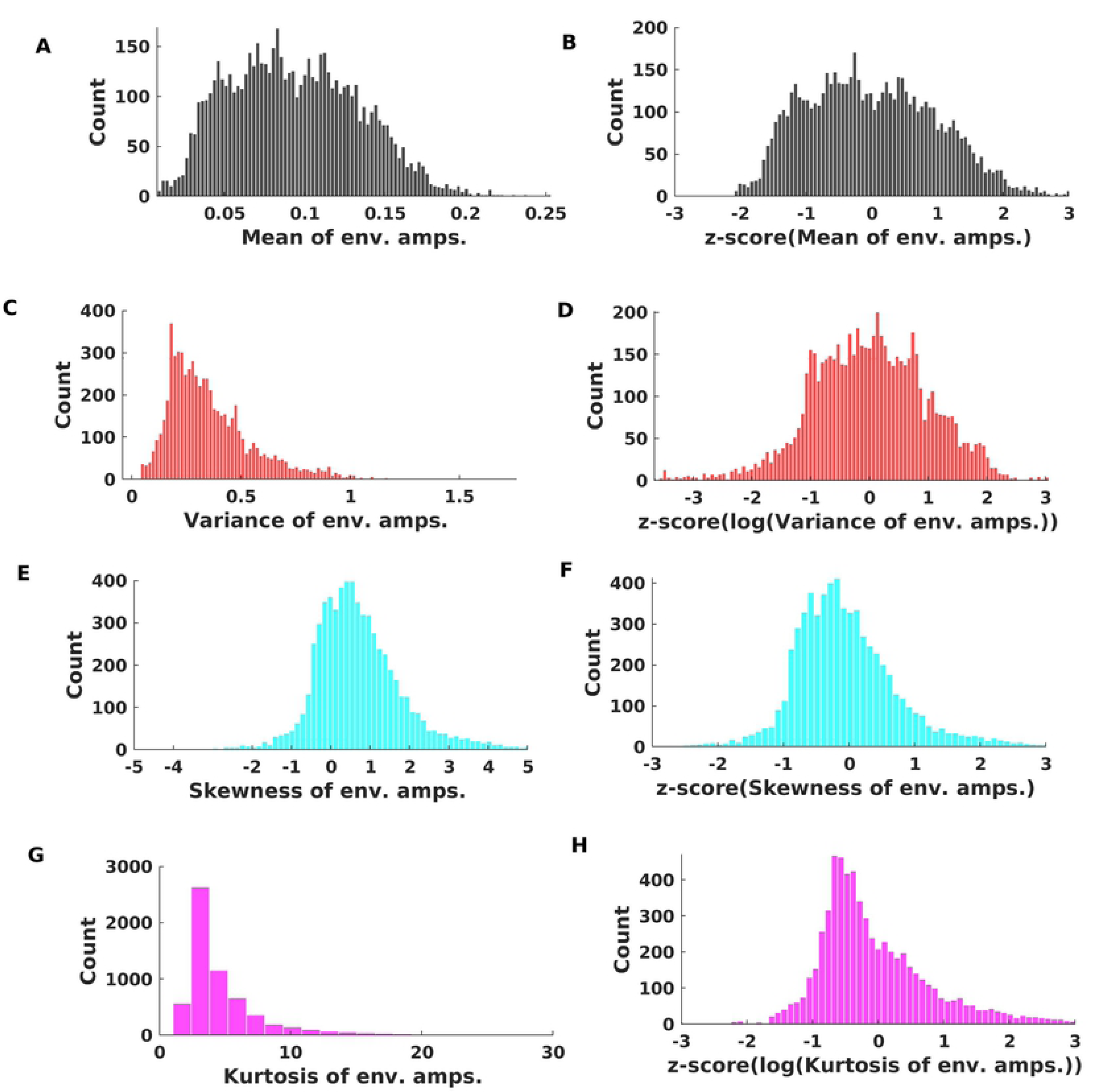
Distribution of statistical parameter values for the entire corpus: (A, B) Distribution of mean of envelope amplitudes. The median, 5th and 95th centile of the distribution were 0.09, 0.034 and 0.16 respectively. (B) Mean of envelope amplitudes after z-scoring. (C) Distribution of variances of envelope amplitudes. The median, 5th and 95th centile were 0.31, 0.13 and 0.77 respectively. (D) Variances of envelope amplitudes after log transformation and z-scoring (E) Distribution of skew of envelope amplitudes. The median, 5th and 95th centile were 0.56, -0.77 and 3.0 respectively. (F) Skew of envelope amplitudes, after z-scoring. (G) Distribution of kurtosis of envelope amplitudes. The median, 5th and 95th centile were 3.79, 2.13 and 17.4 respectively. (H) kurtosis of envelope amplitudes after log transformation and z-scoring.

### 2.1 Principal Components of the Marginal Statistics of Sound Textures

The results of the PCA on the marginal statistics are shown in Fig 3. A key indicator of whether PCA is a useful and appropriate tool to identify major underlying trends and patterns in the data is whether the first few principal components capture (“explain”) a large proportion of the variability between samples. The proportion of variance explained by the first few principal components (PCs) is shown in Fig 3A. Perhaps surprisingly, the first two principal components are sufficient to capture about 65% of the variance of the 128 marginal parameters for our corpus of 200 acoustically very diverse samples of environmental sounds. Fig 3B shows the distribution of our sound corpus over the “marginal space” spanned by the first two principal components, and Fig 3C and 3D show the “shapes” of the first and second PCs for the marginals. When inspecting the heatmap plots of these PCs, it is worth remembering that, because the parameters were z-scored prior to PCA, the units of the color scale are standard deviations above or below the mean parameter values for the entire corpus. The first PC (Fig 3C) is characterized by low means but large variances and skews, with perhaps slightly above average kurtosis, and these trends apply more or less uniformly across all cochlear frequency channels. Consequently, PC1 will discriminate sounds that are “sparse”, with silent periods punctuated by occasional bursts of sound which drive the large variance and skew in the envelopes. The low mean envelope values compensate for the large variance and positive skew: as all sound samples were normalized for RMS power, sounds that are characterized by bursts with positive skew in their envelope amplitudes must have a relatively lower mean envelope “baseline”).

**Fig 3.**
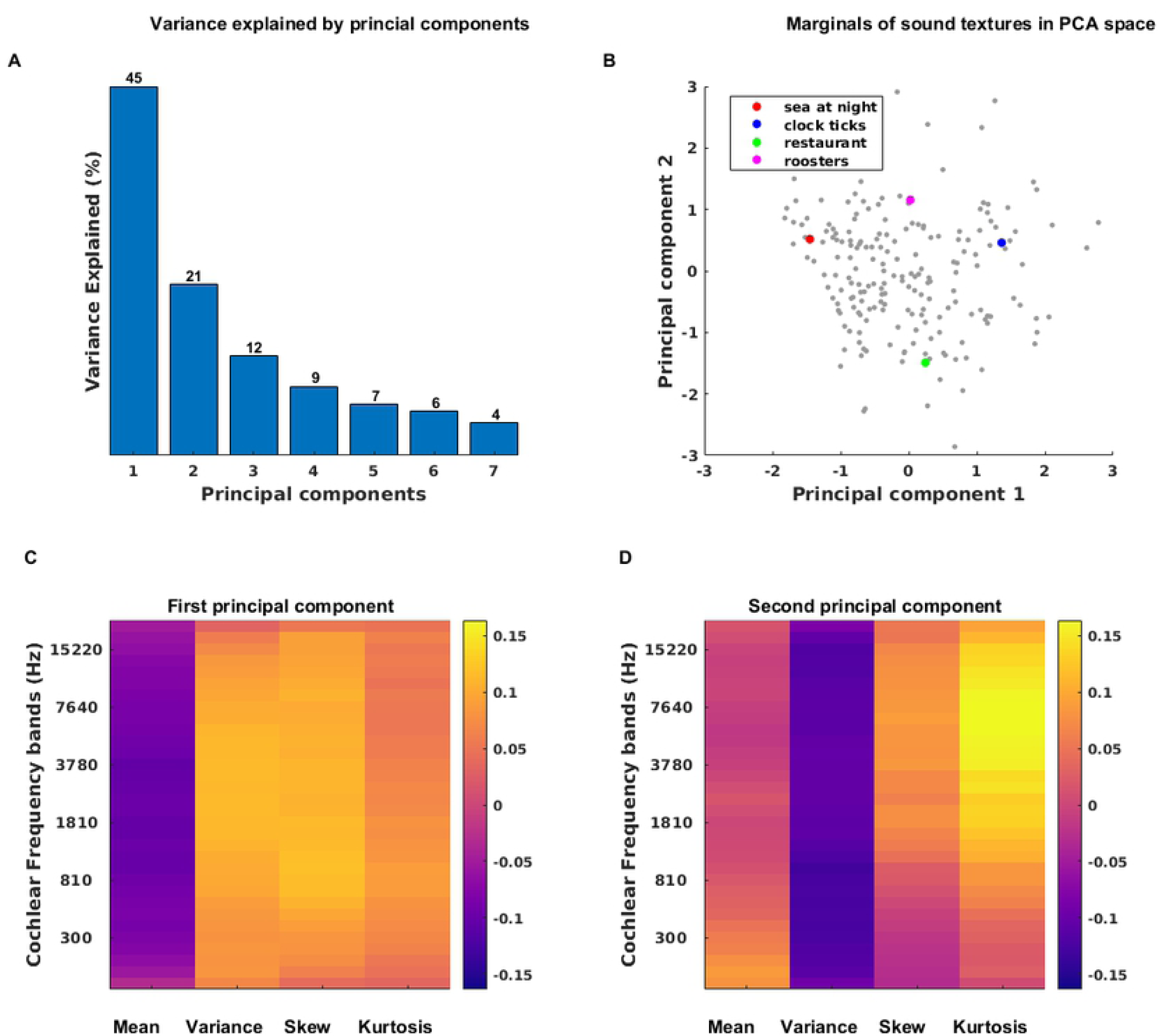
Principal Components of the Marginal Parameters. (A) Percent variance explained by the first 7 PCs of the marginal parameters. The first two PCs capture 65% of the variance explained. (B) Distribution of the sounds in our corpus along the first two PCs of the marginals. Four example sound textures examined further in Fig 4 are highlighted in color. (C, D) Shape of the first and second PCs of the marginals, respectively. The 1st PC distinguishes textures of relatively low mean and high variance, skew and kurtosis from textures for which the reverse is true. The 2nd PC has mean and skew values that are near zero, and thus mostly distinguishes textures with low variance and but high kurtosis, particularly for frequency bands above 800 Hz, from sounds with the opposite feature combination.

To illustrate that the first PC of marginals distinguishes sound textures along a “sparseness” dimension, we examine the marginal statistics of two example sounds from our corpus, “sea at night” and “clock ticks”, in Fig 4. These sounds in the PCA space are highlighted by red and green dots respectively in Fig 3B, and they were chosen to be approximately at opposite ends of the distribution along PC1 but with nearly identical PC2 values. “Sea at night” has lower PC1 values in contrast to “clock ticks”. From Fig 3C, we expect that “clock ticks” should have on average lower envelope means, but higher envelope variance, skew and kurtosis, than “sea at night”. The panels in Fig 4 confirm this. “Clock ticks”, a texture of the sound of multiple clockworks - and brief silences between the ticks - is also a much “sparser” sound than “sea at night”, which features rolling wave and wind sounds that produce continuously elevated sound pressure levels. Others before us have remarked that marginal moments can capture the sparseness of natural stimuli [1, 7, 8], but it is interesting to note from Fig 3A that the first PC of the marginal distributions accounts for almost half of the variability observed in our corpus of natural sound textures, suggesting that the “sparseness” it captures is indeed a major discriminating feature of environmental sounds.

**Fig 4.**
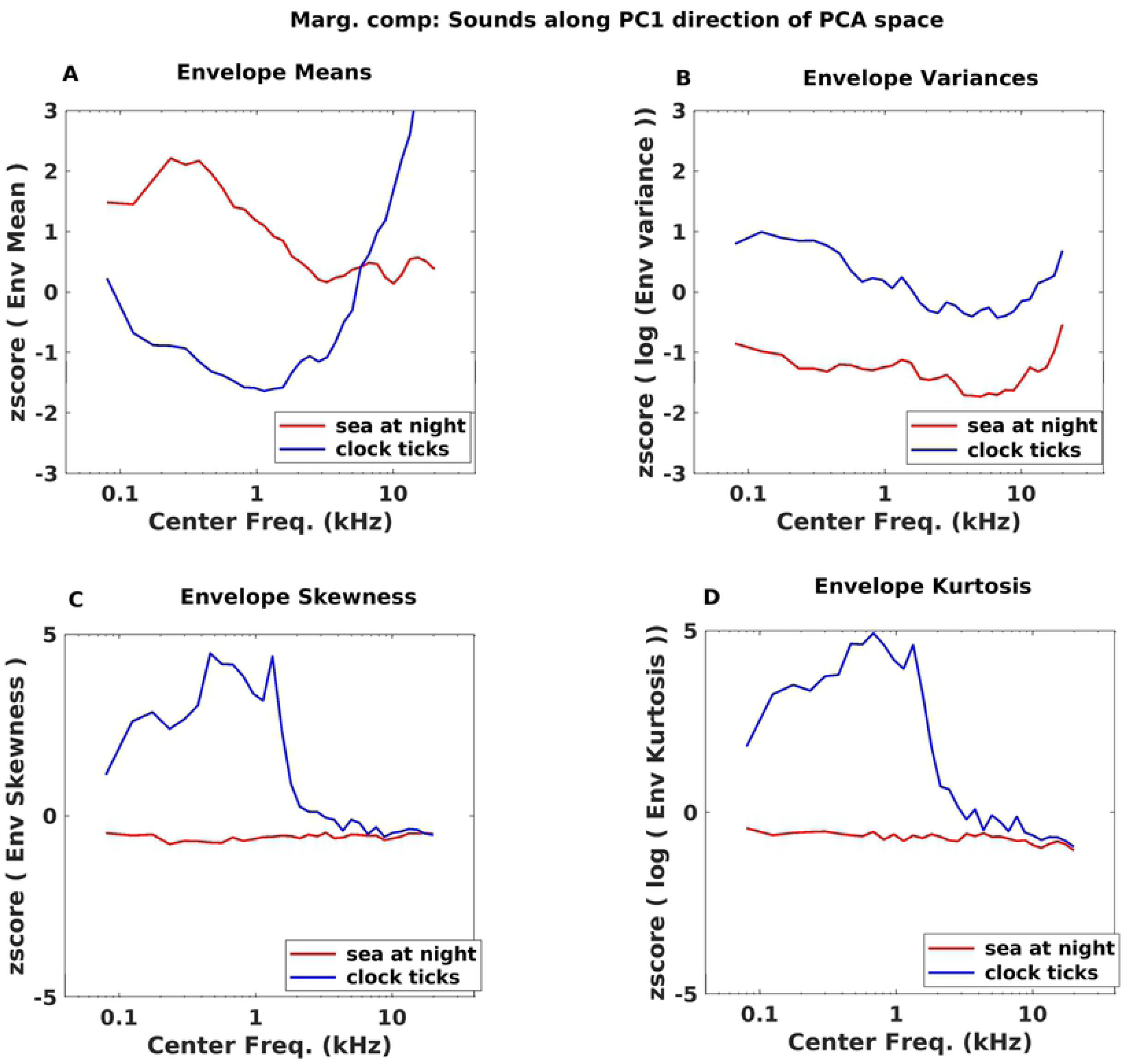
Comparison between the envelope statistics of “sea at night” from one end of PC1 dimension and “clock ticks” from the other end. (A) “Sea at night” has higher envelope mean than “clock ticks” as it is in the lower end of PC1 dimension(B, C, D) Envelope of “clock ticks” with high variance, skewness and kurtosis than “sea at night” for frequencies above 800Hz.

The first PC in Fig 3C can reasonably be interpreted as capturing the sparseness of sounds, but does the second PC shown in Fig 3D also lend itself to an intuitive interpretation? The 2nd PC is characterized by envelope mean z-scores near zero, relatively small (negative) values for variance, but large values for skew and particularly for kurtosis, the latter with some high-frequency bias. To interpret this result, consider that variance, skew and kurtosis all measure excursions from the mean, but skew and kurtosis, as higher order moments, are “more sensitive” to such excursions, growing with the third and fourth powers of the deviation from the mean respectively, rather than just the square. Thus, an envelope distribution with a large kurtosis but a small variance will have a particularly long, thin “tail”, meaning that sound amplitudes can shoot up to very large values relatively frequently, but will not spend much time at “middling” amplitude levels, while for a texture with relatively larger variance and smaller kurtosis, the converse is true. We would therefore expect sounds with large marginal PC2 scores to be not just sparse, but “bursty”, exhibiting intermittent bouts of very high sound energy and fluctuating quite wildly between loud and quiet, but relatively little in between, unlike textures with low PC2 scores which would exhibit comparatively “less extreme” amplitude fluctuations. In PC2, large kurtosis goes hand in hand with positive skew. This is likely attributable to the fact that sound envelope amplitudes cannot be negative, and the large amplitude excursions of “bursty” sounds with high kurtosis are therefore bound to be positively skewed. Thus, PC2 appears to rank sound textures on how “bursty” they are. In Fig 5 we illustrate the marginal statistics for two sounds chosen to vary systematically along PC2, but have approximately the same values for PC1: “restaurants”, and “roosters”. Fig 3C highlights the coordinates of these two sounds in marginal PC space with a green and a magenta spot respectively, and shows that “roosters” has a much larger PC2 value than “restaurants’. As can be seen in Fig 5, the two sounds exhibit the expected trends, with “roosters” having on average smaller variance but greater kurtosis than “restaurants”. Both sounds are of average “sparseness”, but while in “restaurants” there is a variety of background sound events of differing levels (voices, cutlery sounds, footsteps, etc), “roosters” jumps wildly between periods of relative quiet and moments of loud and forceful crowing, making it the substantially more “bursty” sound of the two.

**Fig 5.**
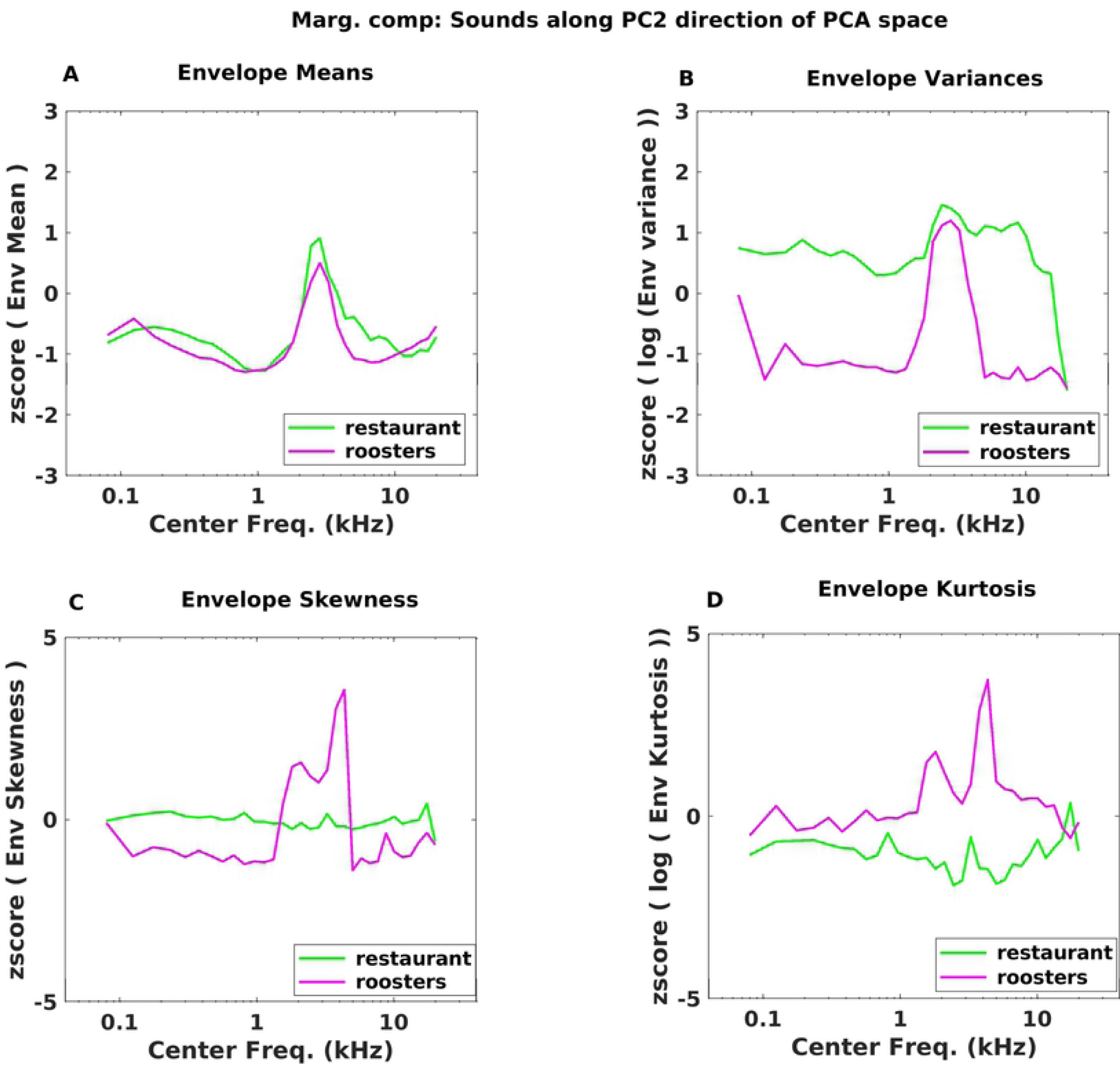
Comparison of “restaurant ambience” and “roosters” across PC2 dimension of Marginal statistics. (A) “roosters” has a lower mean for cochlear envelope than “restaurant”(B) “roosters” has a higher variance cochlear envelope than “restaurant”. PC2 in Fig 3D indicates that as we move along the PC2, sounds should have opposing trends in mean and variance values of their cochlear envelopes. (C, D) As we move in the PC2 direction “skew” and “kurtosis” should be higher. “roosters” has higher skew and kurtosis than “restaurant”.

In summary, the first two PCs of the marginals of our corpus of sound textures between them account for two thirds of the variance (45% and 21% respectively, see Fig 3A) in envelope marginals across the corpus, indicating that the marginal statistics in environmental sound textures are highly redundant. We also observed that the two first PCs lend themselves to an intuitive interpretation, capturing features that can be described as the “sparseness” or the “burstiness” respectively of the sounds.

### Principal Components of the Cochlear Correlations of Sound Textures

The distributions of the cochlear correlations between the envelope amplitudes for different cochlear frequency bands computed for the corpus and pooled over all frequency bands, are shown in Figure 6A. The distribution shows a number of interesting features. Firstly, anticorrelations (that is, negative correlation coefficients) are extremely rare. That is perhaps unsurprising given that positive correlations between frequency bands arise easily whenever a broadband source modulates activity simultaneously in multiple adjacent frequency channels, but physical mechanisms that would lead to anticorrelated sound envelopes in different frequency bands are hard to envisage. Secondly relatively large correlations (R>0.5) are somewhat more common than smaller ones (R<0.5), although the full positive range of correlation coefficients is very well represented. The median R value was 0.5536, and the 5th and 95th centile values were 0.0052 and 0.9163 respectively. Figure 6B shows the distribution of correlations after pre-processing for the PCA via z-scoring to achieve a more symmetric distribution.

**Fig 6.**
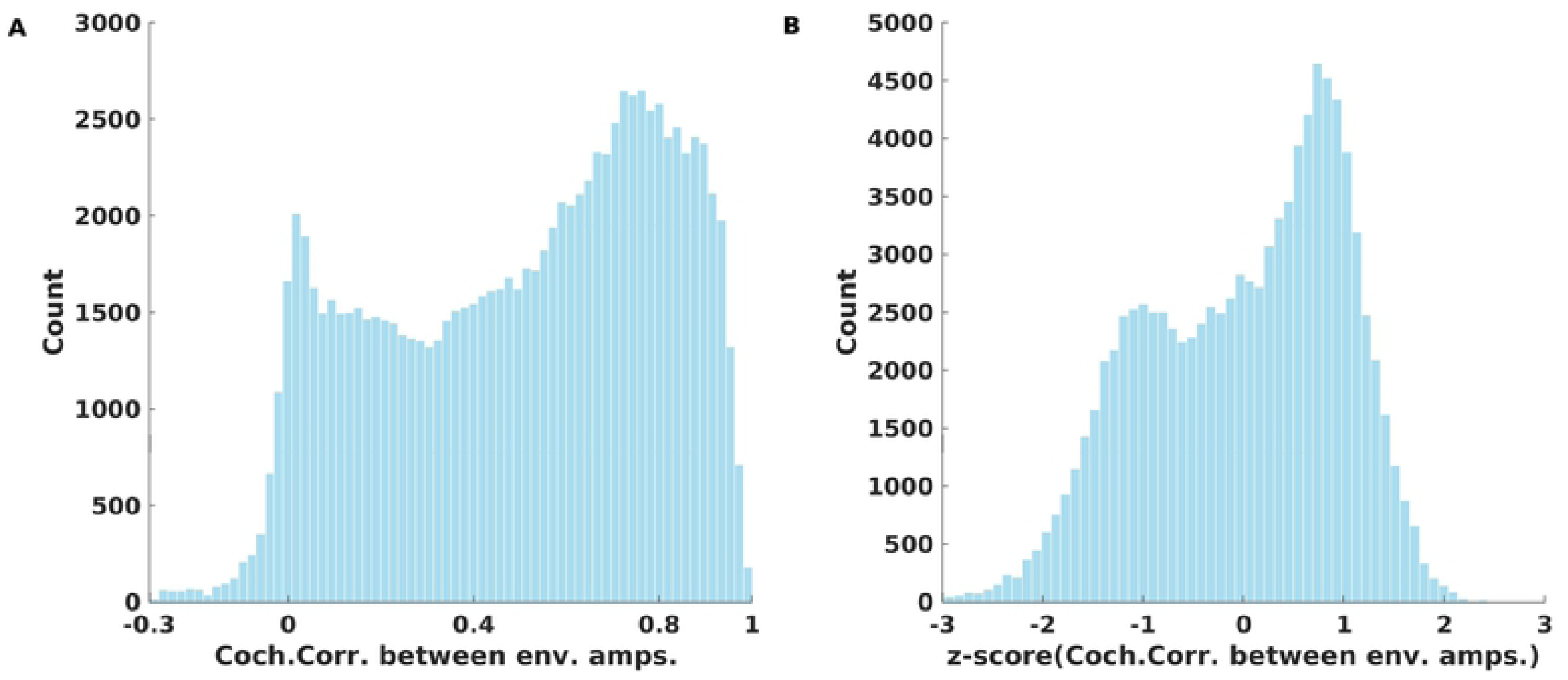
Distribution of cochlear correlation parameter values for the entire corpus: (A) Distribution of cochlear correlations between the envelope amplitudes of different cochlear frequency bands. The median, 5th and 95th centile of the distribution were 0.55, 0.0052 and 0.91 respectively. (B) Distribution of cochlear correlations after z-scoring.

The results of PCA on the correlation statistics are depicted in Fig 7. In Fig 7A we can see that the first PC accounts for a remarkably high proportion of the variance, with 79%. The second PC, in comparison, captures a comparatively modest 11% and the percent variance explained by the remaining components is in the single digits. Despite the great diversity of the corpus and the large number of correlation parameters, 90% of the variability in correlation parameters can be accounted for by the first two principal components only. Fig 7C and 7D show the “shapes” of the first and second PCs for the correlation statistics. The units of the color scale in the heatmaps (Fig 7C and 7D) are once again standard deviations of the correlation values for the entire corpus. Only the upper triangular matrix of the PCs is shown, as these are symmetric correlation matrices.

**Fig 7.**
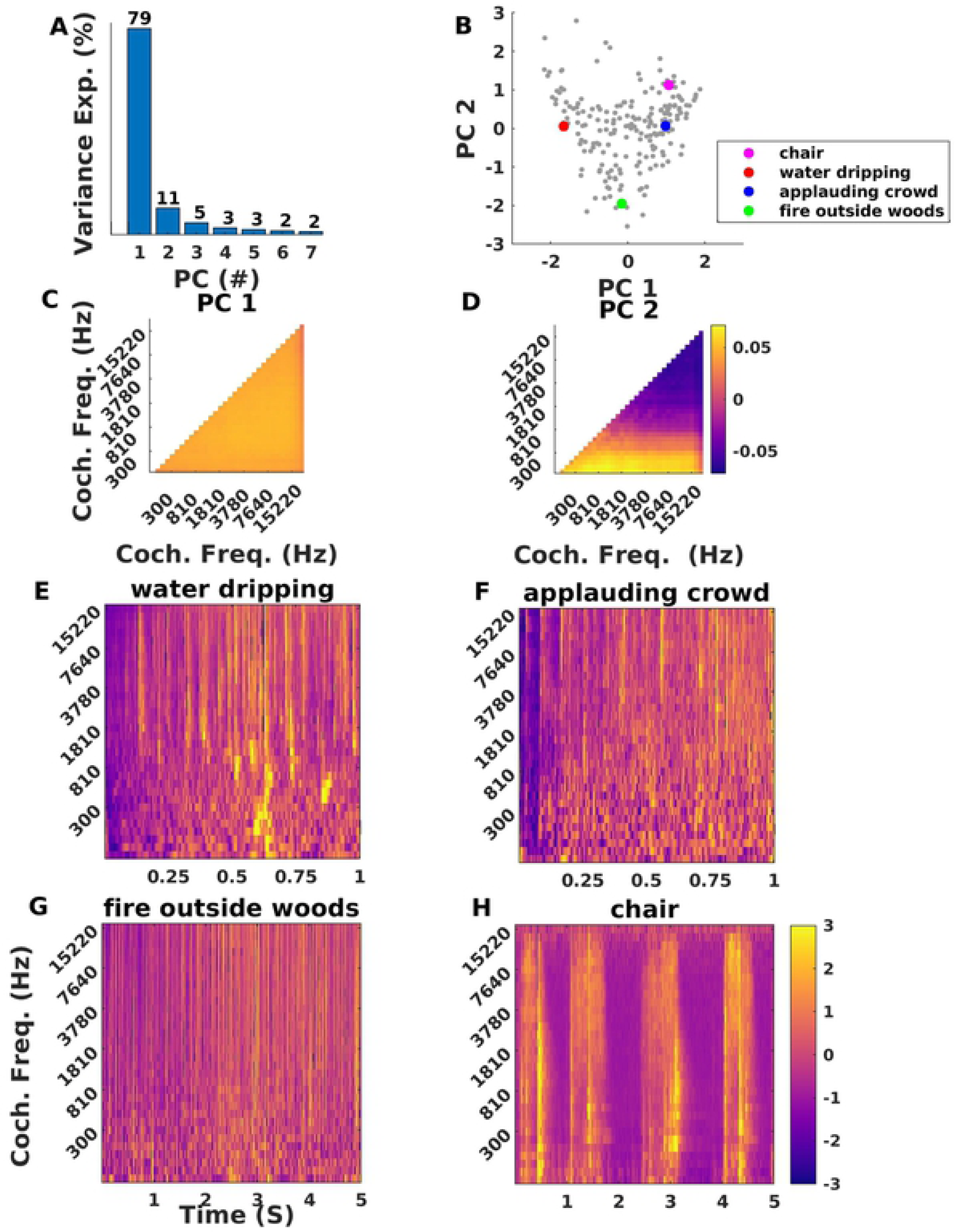
Principal Components of Cochlear Correlation Parameters. (A) The first PC alone captures 79% of variance, whereas the second PC captures merely 11% and higher PCs capture only very small proportions of the variance. (B) Coordinates of all sound textures in our corpus. Coloured dots represent four example sounds examined further in panels E-H. (C) PC1 shows elevated correlation across all pairs of cochlear frequencies, and thus distinguishes “highly correlated” from “poorly correlated” sounds. (D) PC2 captures whether correlations are more pronounced among low or high frequencies. (E-H) normalized cochleagrams for the four sound texture examples highlighted in (B) by colored dots.

The first PC (Fig 7C) is essentially completely “flat”, and it will therefore distinguish textures for which envelope amplitudes are highly correlated between frequency bands from those that are poorly correlated, irrespective of frequency. High correlations among frequencies in sound textures typically arise if many broad-band clicks or noise-bursts contribute to the texture, as these will create synchronized, abrupt changes in sound level across many frequency bands. Thus, an applauding crowd, or pouring gravel onto a hard surface would generate highly correlated sound textures. Examples of less correlated textures are generated from sources that are more narrow band and which become active independently. The sound of running water is a typical example. In running water, much of the sound is created from the excitation of small air bubbles. Each bubble is a resonator with a more or less narrow band resonance that depends on the bubble’s size, and different sized bubbles may burst or become otherwise excited at different times, creating sound patterns that are poorly correlated across frequency.

Indeed, the first PC is very good at discriminating “applause”-like sounds from “water”-like sounds, as can be appreciated from Fig 7B, 7E and 7F. Fig 7E and 6F show the cochleagrams for a sample of dripping water sounds and the sound of an applauding crowd respectively. These cochleagrams have been normalized or sound level in each band by z-scoring the envelope amplitudes in each band independently. Correlations across cochlear frequency bands are visible as vertical bands in these cochleagrams, and the normalization ensures that such bands are not obscured by overall sound level differences in different sound frequency bands. The “dripping water” sound shown in Fig 7E is relatively weakly correlated, as can be seen from its low PC1 coordinate in Fig 7B (red dot), and while there are clear horizontal stripes in the high frequency part of its cochleagram, there are also many prominent narrow band features, particularly in the lower frequencies. In contrast, the “applauding crowds” sound in Fig 7F has a high PC1 correlation coordinate (Fig 7B, blue dot), and a lot of prominent vertical striping throughout its cochleagram.

The second PC of the correlation parameters captures whether correlations are more prominent in low or high frequency bands (see Fig 7D). Normalized cochleagrams of sound textures with very different PC2 coordinates are shown in Fig 7G and 7H. The sound texture “fire outside woods” (Fig 7G, green dot in Fig 7B) has a strongly negative PC2 coordinate, and indeed, the vertical stripes that are characteristic of envelope correlations are prominent only in the higher frequency bands. These features likely originate from broad band but somewhat high-pitched crackling sounds which come about when small twigs in a fire burst, and which contribute to the characteristic “fire” sound. In contrast, the “dragging chair” sound texture (Fig 7H, cyan dot in Fig 7B has a higher PC2 coordinate and correlations are more or less evenly distributed throughout the frequency bands.

In summary, while the number of possible pairwise correlations between cochlear frequency bands is very large (1024 parameters per sound texture in our study), almost 80% of the variance in these correlation statistics is captured by a PC that simply measures the extent to which amplitude envelopes are correlated regardless of frequency band. A relatively modest additional 11% of variance is explained by a PC that distinguishes sounds with correlations in high frequencies from sounds with correlations in lower frequencies. Like marginals, correlation statistics are therefore very highly redundant.

### Principal Components of the Modulation Power Statistics of Sound Textures

The distributions of the original and transformed (pre-processed) modulation power parameters pooled over the entire corpus are shown in Figure 8. Fig 8A shows that the original modulation power distribution for the sound corpus is highly asymmetric and positively skewed. Its median was 0.0278, and its 5th and 95th centile were 0.0037 and 0.1141 respectively. Log transformation and z-scoring for PCA preprocessing made the distribution more much symmetric (Fig 8B).

**Fig 8.**
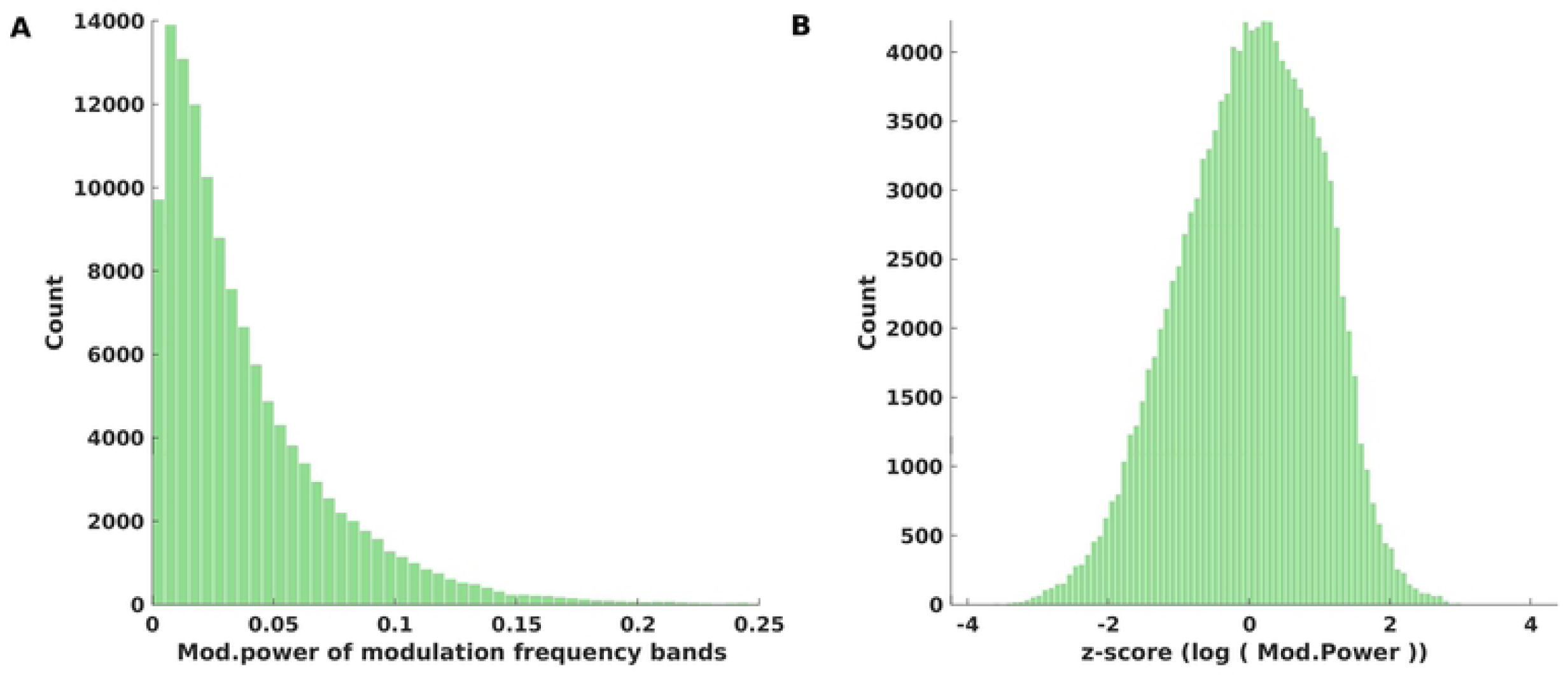
Distribution of modulation power parameter values for the entire corpus: (A) Distribution of modulation power parameters for the entire corpus which are computed over the cochlear envelope amplitudes after modulation filtering. The median, 5th and 95th centile of the distribution were 0.028, 0.0037 and 0.11 respectively. (B) Distribution modulation power parameters after log transformation and z-scoring.

The results of PCA on the modulation power parameters of our sound corpus is shown Fig 9. Analysis of the 640 dimensional modulation power statistics indicates that 73% of variability of our sound corpus are explainable by the first two principal components as shown in Fig 9A. The distribution of our sound corpus along the first two PC dimensions is shown Fig 9B, while Fig 9C and 9D depict the “shapes” of first and second PCs respectively. PC1, which captures 48% of the variance in modulation parameters, discriminates sounds which are modulated at fast modulation frequencies (greater than 60 Hz) from those that are slowly modulated, and this again holds in a very similar manner across all cochlear frequency bands (Fig 9C). Meanwhile, PC2 (shown in Fig 9D) accounts for 25% of the variance and is sensitive to the extent to which sound textures exhibit amplitude modulations at “middling” modulation frequencies of around 30-100 Hz.

**Fig 9.**
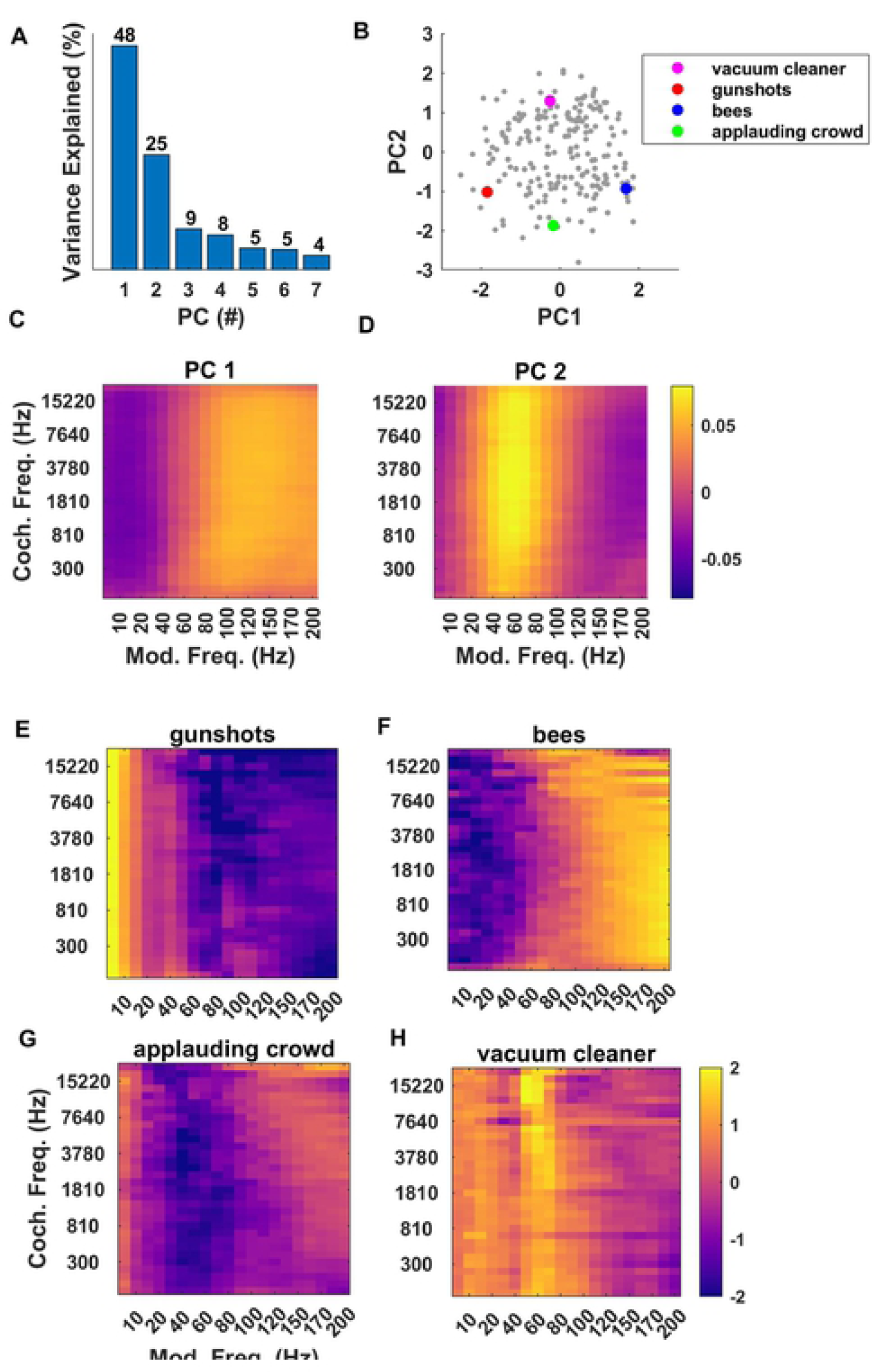
Principal Components of the Modulation Parameters. (A) Proportion variance explained by the first seven PCs of modulation power parameters. (B) Distribution of sound textures from our corpus along the first two PC coordinates for modulation power. The first two PCs capture ∼73% of the variance between them. (C, D) Shape of first and second PC respectively. PC1 discriminates sounds “slowly” from “rapidly” modulating sounds, with a boundary near 60 Hz for all cochlear frequencies. PC2 discriminates sounds with prominent modulations in a “mid range” (near 60 Hz) from sounds lacking such modulations. (E) Modulation spectrum of sound texture sample “gunshots”, showing prominent modulation at low rates. (F) Modulation spectrum for “bees”. High modulation frequencies (> ∼80 Hz) dominate. (G, H) The modulation spectrum for “applauding crowd” shows a relative dearth of modulations near 60 Hz, while that for “vacuum cleaner” shows prominent ∼60 Hz modulations.

We again illustrate these dimensions with examples chosen from the corpus which span either the first or the second PC axes, and which are highlighted in Fig 9B with colored dots. Thus “gunshots” (red dot in Fig 9B, z-scored modulation spectra shown in Fig 9E) lies at the low end of PC1, and the texture is dominated by modulation frequencies of typically less than 10 Hz, while “bees” (blue dot in Fig 9B, z-scored modulation spectra shown in Fig 9F) is dominated by high modulation frequencies, typically above 100 Hz. The causes of the different amplitude modulation rates for these two examples are intuitive: bees beat their wings at much faster rates than users of firearms typically pull triggers. Meanwhile the texture sample “applauding crowd” (Fig 9G, green dot in Fig 9B) has a PC2 coordinate of approximately -1.8, and its modulation spectrum exhibits the expected dearth of modulations near 60 Hz, while the texture “vacuum cleaner” (Fig 9H, cyan dot in Fig 9B) has a PC2 coordinate of ∼+1.5 and prominent ∼60 Hz modulations.

In summary, just like marginal and correlation parameters, modulation parameters too exhibit a high degree of redundancy, so that almost three quarters (73%) of the variance across the 640 parameters could be captured with just two PC coordinates. And again, the PCs obtained lend themselves to simple interpretations, in this case fast vs slow (for PC1), and with or without much modulation in a mid, ∼60 Hz range (for PC2).

## 3 Discussion

The idea that statistical regularities may govern the types of sensory stimuli we encounter in our environment has a long history, as does the idea that the sensory systems may be adapted to some of these statistical features or regularities [11, 12]. This idea has arguably been much more influential in vision research than in hearing research. For example, an attempt by [13] to explain the centre-surround structure of primary cortex visual receptive fields as nature’s solution to the problem of having to encode the structure of visual scenes in a sparse, and hence energy-efficient, manner, has become enormously influential. (Note, however, that more recent work by [14] proposes an intriguing alternative explanation, namely that cortical receptive field structure not just of visual but also auditory cortical neurons may be optimized to facilitate prediction of future inputs, rather than energy efficiency.)

An early example of work looking for statistical regularities specifically in the auditory modality comes from [15], who already reported over 40 years ago that pitch and amplitude fluctuations over long segments of music and speech streams recorded from the radio exhibited a so-called 1/f distribution. Garcia-Lazaro and colleagues [16] later built on that observation and showed that auditory cortex neurons appear to be tuned to these statistics, in that they respond more strongly and reproducibly to artificial sound streams that follow 1/f distributions than to sounds which fluctuate according to slower (1/f0.5), or faster (1/f2) distributions. This was later shown to be an emergent property of the ascending auditory pathway, as inferior colliculus neurons generally prefer more rapidly fluctuating sounds, and neurons in the medial geniculate exhibited no particular preference for fluctuations that were either faster or slower than 1/f [17]. These studies are conceptually similar to earlier work by Also highly relevant are studies by [7, 18] which had described power-law statistics in amplitude distributions of natural sounds, and reported evidence that midbrain neurons can encode synthetic sounds with higher accuracy (as quantified by mutual information), when these stimuli match the statistical parameters typical of natural sounds. Other noteworthy examples of studies concerned with the distributions of environmental sounds and their relevance to auditory processing include a well-known study by [19], which presented an efficient coding argument alongside an analysis of natural sounds to explain the cochlear frequency tuning characteristics, or a study by [20] which described the low-pass nature of spectral and temporal modulations in natural sounds, in a manner corroborating and extending the findings by [15].

Despite this relatively long history, the literature on natural sound statistics and their relevance to auditory processing and perception has remained relatively thin, perhaps because it is still unclear which of the many statistical parameters that could in theory be devised or applied to the study of natural sounds is most likely to provide highly useful and practical descriptors of natural sounds. In this context, the study by [1] provided a fresh perspective. By being able to generate recognizable, and often highly realistic, morphs of natural sound textures by imposing the statistical parameters they had identified onto white noise, [1] demonstrated that their chosen statistical parameter set comes close to being a set of “sufficient statistics”. The fact that these sets of parameters fully describe many types of natural sound textures also raises the intriguing hypothesis that neurons in the central auditory system may be tuned to statistical parameters similar to those identified in their study. Such tuning could easily explain our perceptual ability to distinguish different sound textures with ease, even though these textures are stochastic signals, and two recordings of the same type of sound texture are essentially guaranteed to be very different sound waves. Identifying a set of statistical parameters that come close to fully characterizing sound textures is therefore a very significant conceptual advance. However, there are issues which make it difficult in practice to build on their work with follow-on psychoacoustic and neurophysiological studies. One such issue is the fact that the number of statistical parameter values used by [1] to characterize and synthesize textures is very large, in the order of 1500 parameters in total for each texture. This parameter explosion arises largely because marginals, correlations and envelope modulation spectra are computed independently for every frequency band. In addition, the range of parameter values that one is likely to encounter in natural or environmental soundscapes has not been described.

Ideally one would wish to build on [1] approach to devise a characterization of the statistical features of natural sound textures which uses a far smaller number of numerical parameters and identify their distributions across the ecological acoustic environment. We do not claim that our analysis presented here has achieved this, but it has nevertheless shown that this may be possible in principle, given the enormous redundancy of the statistical parameters we have identified through our PCA of a corpus of environmental texture recordings which we have analysed. Indeed, just two PC coefficients for marginals captured 66% of the variance of a 128 dimensional parameter space.

Similarly, the first two PC coefficients of cochlear correlations cover 90% of 1024 dimensional parameter space, and the first two PC coefficients of modulation power explain 73% of variance of a 649 dimensional modulation parameter space. Lower-dimensional descriptions of natural sound statistics which nevertheless capture much of the richness of the auditory environment should therefore be possible.

Another noteworthy finding of our PCA analysis is that it illustrates the high degree to which many statistical features tend to co-vary greatly across frequency bands. Thus, the first PC across the marginals showed very little variation in Mean, Variance of Kurtosis as a function of cochlear frequency (Fig 3C), the first PC of cochlear correlations is effectively constant across all pairs of frequency bands (Fig 6C), and the first and second PCs of the modulation parameters too exhibit very little variation as a function of cochlear frequency. This observation is not entirely novel, as [7] had already conducted a filter bank analysis on an ensemble of natural sounds, and reported that temporal lower order statistics for a given sound sample tend to be highly similar across frequency bands. Nevertheless, in combination with the many additional values which we report here, this confirmatory finding is potentially quite useful. Thus, if someone presented us with a “mystery texture sound”, reproduced at a unit RMS amplitude, and asked us to guess what its statistical parameters are likely to be in some particular frequency band, then we would be able to declare with some confidence, firstly, that the particular frequency band probably does not matter, secondly, that its mean envelope amplitude has a 90% chance of falling between ∼0.0338 and 0.1618 with a maximum likelihood value of ∼0.0905 (Fig 2A), the variance of the envelope amplitude has a 90% chance of falling between ∼0.1292 and 0.7763 with a maximum likelihood of ∼0.315 (Fig 2C), its skewness has a 90% chance of falling between ∼-1.8 and +3 with a maximum likelihood of ∼0.6 (Fig 2E), and its kurtosis is 90% likely to fall between ∼1 and 18 (Fig 2G) with a maximum likelihood of ∼5. Similarly, envelopes in any two cochlear frequency channels are a priori more likely than not to be substantially correlated, with an R > 0.55 (Figure 6).

The data presented here can therefore facilitate informed guesses about as yet unknown natural sounds that we may be presented with in the future, and we hope that a better characterization of the statistical features of natural sounds will enable us to start asking better questions about the extent to which expectations derived from these distributions may be “built into” the functional anatomy of our central auditory nervous system.

## Acknowledgments

Funding for this project was provided by the Hong Kong General Research Fund (No. 11100617).We also thank Fei Peng for helpful discussions on drafts of the manuscript

